# High-density theta burst stimulation (hdTBS) at 100 Hz triples the aftereffects of the conventional intermittent TBS

**DOI:** 10.1101/2025.09.15.676379

**Authors:** Aidan Carney, Taylor Scott, Olivea Varlas, Md Mohaiminul Haque, Hieu Nguyen, Yihong Yang, Hanbing Lu

**Affiliations:** Neuroimaging Research Branch, Intramural Research Program, National Institute on Drug Abuse (NIDA), National Institutes of Health (NIH), Baltimore, MD, USA; Electrical and Electronic Engineering, University of Nebraska-Lincoln, Lincoln, Nebraska, USA; School of Electrical Engineering, International University, Ho Chi Minh City, Vietnam; Vietnam National University, Ho Chi Minh City, Vietnam

**Keywords:** Theta burst stimulation (TBS), motor evoked potential (MEP), motor threshold, transcranial magnetic stimulation (TMS), long-term potentiation (LTP)

## Abstract

Slice electrophysiological studies have experimentally demonstrated that theta burst stimulation, consisting of electrical pulses delivered at 10 ms (100 Hz) inter-pulse intervals, optimally induces long-term potentiation in the hippocampus. Inspired by this observation, a novel transcranial magnetic stimulation (TMS) paradigm, 100 Hz high-density theta burst stimulation (100 Hz hdTBS), is presented. This paradigm delivers 6 pulses per burst with an inter-pulse interval of 10 ms – doubling the pulse frequency and total pulse count of the conventional intermittent TBS (iTBS). The effect of this new paradigm was studied in the motor cortex of awake rats using a rat-specific focal TMS coil and a hdTBS stimulator developed in house. Results reveal that 100 Hz hdTBS triples the after-effects of conventional iTBS. In a separate group of animals that received two consecutive iTBS session back-to-back (prolonged iTBS), we observed an inhibitory effect. Since that prolonged iTBS matches the total pulse count of 100 Hz hdTBS but produced opposite after-effects, our results underscore the critical roles of the temporal structure of TMS pulses—not merely the total number of pulses—in driving neuroplasticity. This new paradigm has the potential to significantly enhance therapeutic efficacy if confirmed to be safe and effective in humans.

## Introduction

Intermittent theta burst stimulation (iTBS) achieves statistically significant, though modest antidepressant efficacy comparable to that of 10 Hz repetitive TMS[1–3]. Many efforts have been made to enhance the effects of TBS, for example, changing pulse shape[4] or temporal patterns[5]. The Stanford Neuromodulation Therapy (SNT) paradigm[6], an “accelerated” version of iTBS, achieved a higher remission rate in major depression. SNT applies 10 TMS sessions per day, 1800 iTBS pulses per session. Note however that conventional iTBS, the building block of the SNT paradigm, exhibits limited effect in modulating cortical excitability[7]. There is a need to develop new TBS technology to enhance efficacy while maintaining high time-efficiency, and to support further treatment optimization.

A long-standing view holds that TMS exerts its therapeutic effects through physiological mechanisms known as long-term potentiation (LTP) and long-term depression (LTD)[3,8]. Slice electrophysiological studies in the 1980s by Larson and colleagues demonstrated experimentally that TBS was most efficient in inducing LTP[9,10]. In these studies, bursts of electrical pulses—each consisting of 4 to 6 pulses—were delivered at 100 Hz, with an inter-burst interval of 200 ms (5 Hz) (referred to as classical TBS hereafter). The transient peak power of these pulses was in the milliwatt range. Translating this TBS paradigm to non-invasive brain stimulation using TMS presents several technical challenges due to its high-power requirement (>3kA, 2kV). Additionally, TMS pulses at high frequencies lead to coil overheat. Perhaps due to these technical challenges, the original iTBS paradigm proposed by Huang et al. in 2005 applied triplet pulses at 50 Hz[11]. We hypothesized that a TMS method that closely mirrors the classical TBS paradigm will yield the most pronounced effects.

We previously introduced a high-density TBS (hdTBS) paradigm[12], which delivered six pulses per burst as opposed to only three in conventional iTBS (referred to as 50 Hz hdTBS hereafter). In studies targeting the motor cortex of awake rats, this paradigm produced significantly stronger after-effects than conventional iTBS. The current study has two objectives: (1) to further enhance the hdTBS protocol by increasing the burst frequency to 100 Hz while maintaining a 200 ms inter-burst interval (referred to as 100 Hz hdTBS), bringing it closer to the classical TBS structure; (2) to assess the after-effects of this new method and compare it with existing methods. Experiments were conducted in the motor cortex of awake rats using the focal TMS coil that we have developed[13]. Results demonstrate that 100 Hz hdTBS triples the after-effect of the conventional iTBS, establishing a novel approach that has the potential to drastically enhance the efficacy of TMS therapy.

## Materials and Methods

### Development of 100 Hz hdTBS stimulator

Figure 1A illustrates the iTBS protocol, as previously described[11]. The stimulation consists of 20 pulse trains, each comprising a 2 s stimulation ON followed by 8 s OFF. During the ON period, bursts of TMS pulses, each consisting of 3 pulses with an inter-pulse-interval of 20 ms, were delivered every 200 ms, totaling 600 pulses per session. The 50 Hz hdTBS paradigm that we reported previously is identical to iTBS except that each burst delivers 6 pulses as opposed to only 3 (Fig. 1B)[12], totaling 1200 pulses per session. The 100 Hz hdTBS paradigm delivers 6 pulses per burst with a reduced inter-pulse-interval to 10 ms (Fig. 1C), totaling 1200 pulses per session. Both the 50 and 100 Hz hdTBS preserved the 200 ms inter-burst interval, a parameter demonstrated to be critical for the induction of LTP[14].

**Figure 1.**
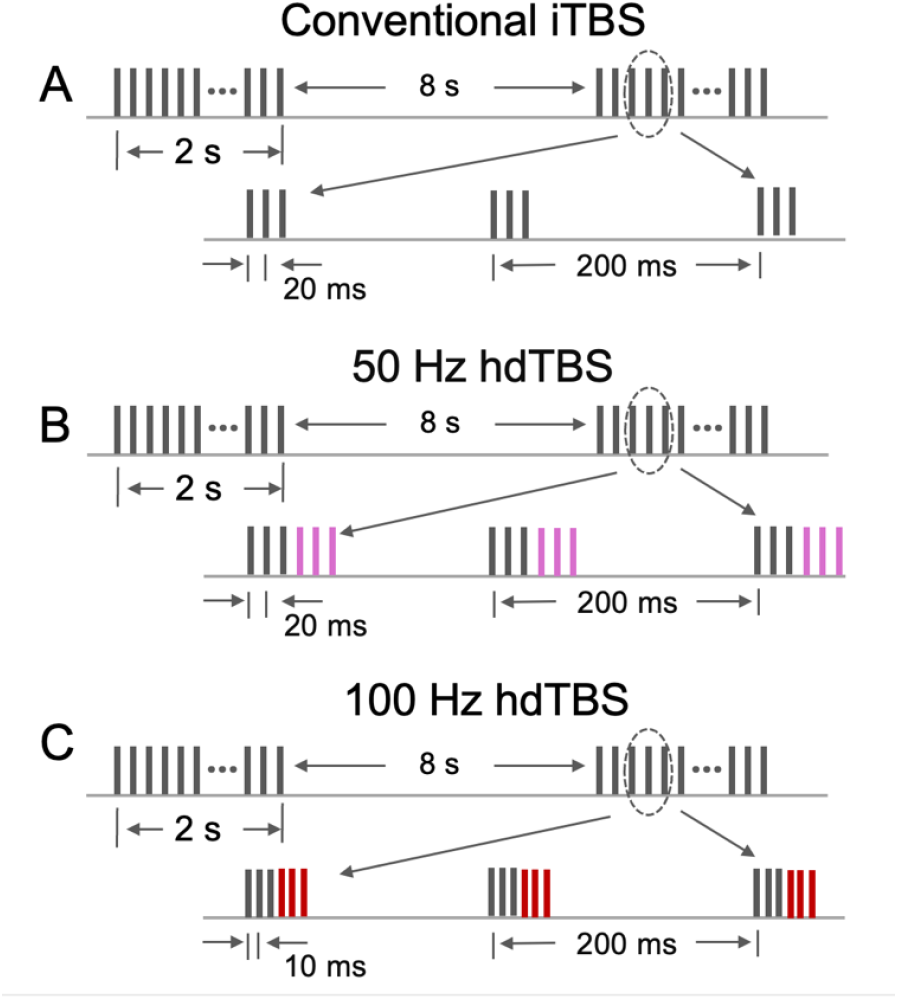
(**A**) Schematic of the conventional iTBS paradigm, which includes 20 cycles of 2 seconds ON and 8 seconds OFF. During each ON period, bursts of pulses are delivered every 200 ms, with each burst consisting of three pulses at 50 Hz, totaling 600 pulses per session. (**B**) The 50 Hz hdTBS protocol follows identical structure as iTBS but increases the number of pulses per burst to 6, resulting in 1200 pulses per session. (**C**) The 100 Hz hdTBS protocol also delivers six pulses per burst but at 100 Hz, while maintaining the 200 ms inter-burst interval.

We in-house developed a stimulator capable of delivering 100 Hz hdTBS pulses (Fig. 2A). The circuit topology was based on our previous report[12], employing high power Insulated Gate Bipolar Transistor technology to switch on and off coil current. Since hdTBS at 100 Hz imposes stringent requirements for power electronics, we implemented a major overhaul in the power supply units, with particular attention to minimizing interference between power supply components.

**Figure 2.**
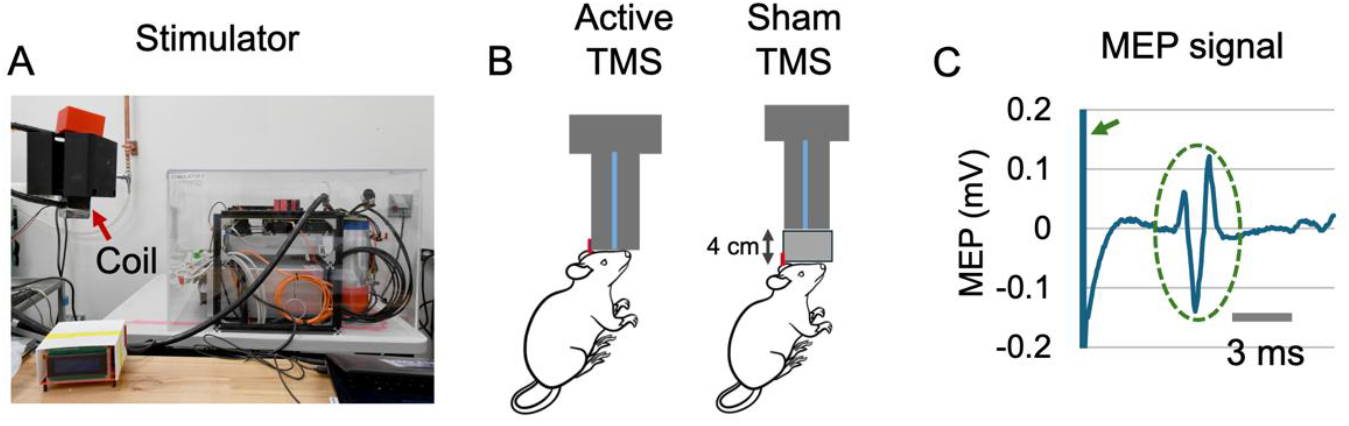
(**A**) The custom-built TMS stimulator prototype. The red arrow highlights the rat-specific stimulation coil. Only high-power electronic components are shown for visual clarity. (**B**) Illustration of experimental set up for active and sham TMS. In the active TMS condition, the rat’s head was positioned in close proximity to the coil surface, and stimulation was delivered at 100% of the motor threshold. In the sham condition, a 4 cm-high plastic spacer was placed between the coil and the rat’s head, and stimulation intensity was reduced to 60% of the motor threshold, the induced E field at 4 cm below the coil surface was effectively zero. (**C**) Representative raw MEP trace. The green arrow marks the TMS-induced artifact, lasting approximately 3 ms, followed by the MEP response, indicated by the green oval.

### Animal experiments

Experiments were performed on adult Sprague-Dawley rats (N = 23). We employed a within-subject design: each rat received one active and one sham TBS session separated by 1 week; the order of active and sham TBS session was counter-balanced within each experimental group.

Animals were randomly assigned to one of three experimental groups: 100 Hz hdTBS (N = 8), conventional iTBS (N = 8) and prolonged iTBS (pro-iTBS; N = 7)[15]. The pro-iTBS protocol consisted of two consecutive conventional iTBS sessions, delivering 1200 pulses per session—matching the pulse count of the 100 Hz hdTBS protocol. This design permits the dissociation of stimulation effects attributable to the total pulse count versus the temporal patterns of the TMS pulses.

Rats underwent two surgical procedures: (1) implantation of microelectrodes into the hindlimb muscles for recording motor-evoked potentials (MEPs), and (2) headpost implantation to accurately align the motor cortex with the hotspot of a custom-built, rat-specific TMS coil. All procedures were approved by the Animal Care and Use Committee of National Institute on Drug Abuse (NIDA), National Institutes of Health (NIH).

### Electrode implantation for MEP recording

We implanted microwire electrodes via a “back-mount” procedure to longitudinally record the MEP signal in awake rats[12,16]. The electrodes were made of soft 7-strand stainless steel microwires (0.025 mm in diameter, A-M systems, Washington, USA), cut into 13 cm in length. The insulation coating on one end of the microwires was removed (about 3 mm in length) before being press-connected to a female socket (Model E363/0, P1 Technology, USA). The sockets were then organized and embedded into an electrode pedestal (Model: MS363, P1 Technology, USA). To ensure the stability of the sockets and microwires, the pedestal was attached to a circular Marlex mesh secured with dental cement. The insulation coating at the other end of microwires was also carefully removed, which were implanted into the biceps femoris muscle and the gastrocnemius muscle of the left hindlimb. These exposed segments served as active electrodes to detect electromyography (EMG) signal in the implanted muscles; one surface EMG pad was attached to rat tail serving as the ground electrode. The two active electrodes and the ground electrode were interfaced to a BIOPAC EMG recording system (BIOPAC Systems Inc, CA, USA). The EMG signal was amplified by a factor of 2000, band-pass filtered between 100 and 5000 Hz, and sampled at 10,000 Hz.

### Headpost Implantation and Design

To ensure precise targeting of the hindlimb motor cortex (M1), a custom-designed plastic headpost was surgically implanted onto the rat’s skull. The headpost features an L-shape with a flat horizontal extension measuring 1 cm in length and a vertical block standing 1 cm tall. The headpost was secured to the skull using dental cement (C&B Metabond Quick Adhesive Cement System, Parkell, USA) and placed such that its anterior edge was 1.8 mm posterior to Bregma. This specific design and placement of the headpost facilitated consistent and accurate localization of the coil hotspot over M1 (AP −1.8 mm, ML 1.5 mm) during stimulation, thereby enabling reproducible coil positioning across experimental sessions.

### Habituation and Motor Thresholds

To minimize stress and ensure cooperation during TMS administration, rats were habituated to the TMS environment prior to receiving stimulation. Habituation was conducted over 5 days. The rat was held under a dummy coil for 2 sessions of 3 minutes (the length of a typical TMS administration session) while single-pulse TMS was running. Rats did not receive any actual stimulation during habituation. Rats slowly acclimated to the sensation of being held up to the coil, including being restrained by experimenters with their ears covered, and having their heads touching the coil. At the end of habituation, rat motor thresholds were determined. Motor threshold was considered the power level required to elicit a left hindlimb response 50% of the time when a single TMS pulse was delivered to the right hindlimb M1.

### TMS experiment and MEP recording

We recorded MEP signal immediately prior to active or sham TBS administration, referred to as the pre-TBS baseline. Ten single TMS pulses were delivered at 100% of the motor threshold, spaced 8–10 seconds apart. The rat’s ears were covered during TMS administration to minimize the effects of the acoustic noises from the TMS coil, which effectively prevented startle responses commonly observed when ears were left uncovered.

Following active or sham TBS administration (see below), we recorded MEP signal at 5, 10, 15, 20, 25, and 35 minutes post-TBS using the same procedures described above (Fig. 3A).

**Figure 3.**
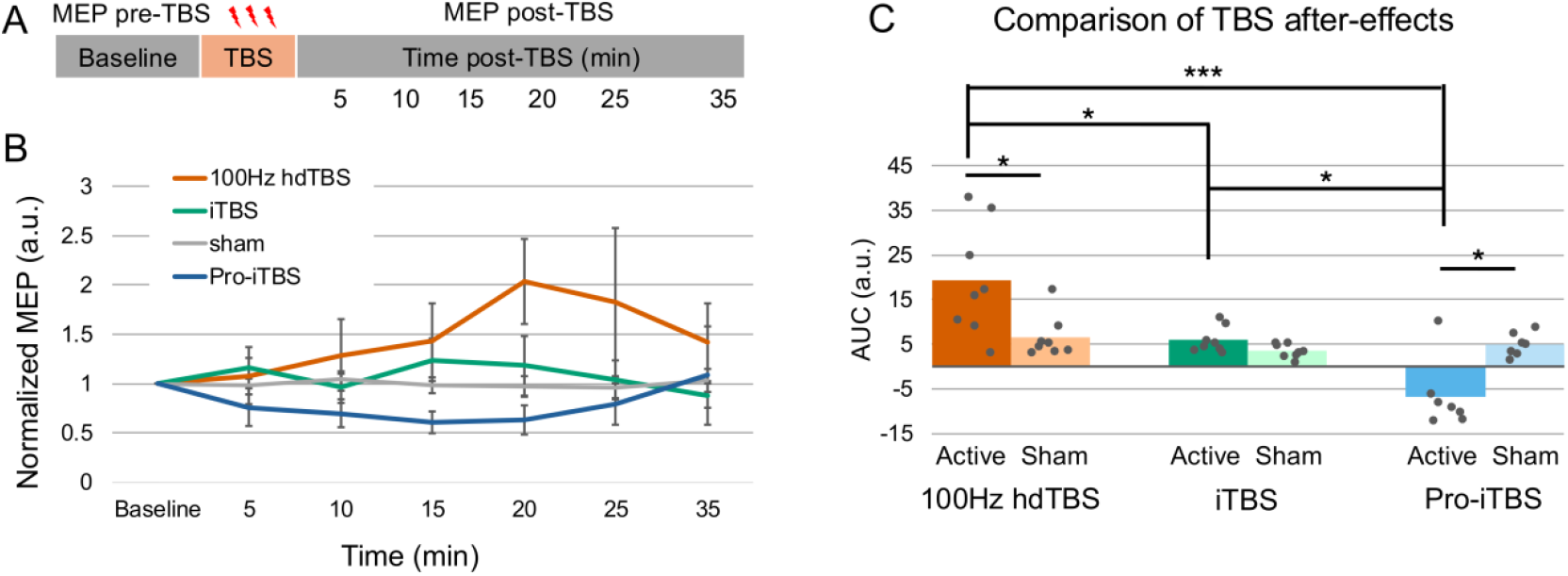
(**A**) Experimental timeline: MEP signal was recorded prior to stimulation and up to 35 minutes following TBS administration. (**B**) Averaged normalized MEP responses for each TMS condition. (**C**) Comparison of AUC values of the MEP responses. Notably, prolonged iTBS, which involves two back-to-back conventional iTBS sessions to match the 1200 pulses per session used in hdTBS protocols, produced a reduction in MEP amplitude, whereas 100 Hz hdTBS led to increased MEP signals despite all delivering the same total number of pulses. Abbreviation: AUC, area-under-curve; pro-iTBS, prolonged iTBS; a.u., arbitrary unit. *, P<0.05; ***, p<0.0005.

For active TBS, the TMS power level was set to 100% of motor threshold, and the rat head was placed in close proximity to coil surface. For sham TBS, TMS power was set to 60% of motor threshold, and a plastic spacer 4cm thick was placed under the coil. The rat head was placed in close proximity to the surface of the plastic block (Fig. 2B) during sham administration. The acoustic noise level was similar under these two conditions. Our measurement confirmed that the electrical (E) field produced by our focal TMS coil was effectively zero at 4 cm below the coil surface. The acoustic noise level was similar under these two conditions. The rat-specific coil was cooled with dry ice and there was an interval of 75 minutes between two TBS sessions to prevent it from being overheated.

### Statistical Analysis

TMS pulse induces characteristic artifacts of large amplitudes lasting for about 3 ms followed by multi-phasic EMG signal that last for about 4 ms (Fig. 2C). MEP data was extracted based on the onset of the artifacts using custom Matlab scripts. Both the peak-to-peak amplitude and area-under-curve (AUC) of the MEP waveforms were calculated. As expected, no significant difference in trend was observed between AUC and amplitude data across time points for each rat. For each rat, pre-TBS baseline MEP signals were averaged (MEP_base_); post-TBS MEP signals at each time point were also averaged (MEP_i_, i=5, 10, 15, 20, 25 and 35), which was normalized to pre-TBS baseline (MEP_i_/MEP_base_). Since MEP amplitude and AUC data were non-normally distributed, we analyzed the data using the multi-factorial Aligned Rank Transform Analysis of Variance (ART-ANOVA), a non-parametric method. TBS TYPES (100 Hz hdTBS active and sham, iTBS active and sham, pro-iTBS active and sham) and TIME (Baseline, post-TBS times) were considered as the two factors for analysis. AUC data was analyzed using the same ART ANOVA method, but considering only one factor, (TBS TYPES). Further post-hoc analysis was performed to determine pairwise comparisons of the above factors, correcting for multiple comparisons using the Benjamini-Hochberg procedure. ART ANOVA and subsequent post-hoc analysis (ART-C) was performed using the ‘ARTool’ package in R[17].

## Results

All rats tolerated the TMS administration well. We did not observe any signs of seizure during or after stimulation, and there were no noticeable changes in daily behaviors such as eating, drinking, grooming, rearing, etc.

Figure 2C shows a representative MEP recording. It features artifacts of large amplitude lasting for about 3 ms (green arrow), followed by multiphasic EMG signal (green oval). We recorded MEP signal at 7 time points: pre-TBS baseline, 5, 10, 15, 20, 25 and 35 minutes post-TBS (Fig. 3A). Data from individual animals were normalized to their pre-TBS baselines, and are presented in Supplemental Fig. 1. Figure 3B presents the average MEP signals across time. To quantify overall response, we calculated AUC for each animal’s post-TBS MEP signal.

The resulting AUC values were: 5.9 ± 2.9 for conventional iTBS, 19.3 ± 12.6 for 100 Hz hdTBS, and –6.8 ± 7.8 for prolonged-iTBS. Statistical analysis was performed using ART-ANOVA, a non-parametric method. Results are summarized in Fig. 3C. There were significant interaction effects between TBS TYPES and TIME (F[30, 250.8] = 3.54, *p* = 2.09×10^−8^). There was significant MAIN effect of TBS TYPES in AUC values (F[5, 38] = 7.0, *p* = 9.95×10^−5^). Post-hoc analysis reveals the AUC values in the 100 Hz hdTBS group were significantly higher than that of the iTBS group (*p* = 0.04); AUC values in the pro-iTBS groups were significantly lower than the other 2 groups (*p* = 2.9×10^−5^ for 100 Hz hdTBS vs. pro-iTBS; *p* = 1.1×10^−2^ for iTBS vs. pro-iTBS). Detailed statistical results are summarized in Fig. 3C and in Supplemental Table 1.

## Discussion

In the present study, we have proposed, developed, and validated a 100 Hz hdTBS paradigm using a rat-specific focal TMS coil in awake animals. Using MEP as the readout[11,18–20], we compared the after-effects induced by this new paradigm with those of conventional iTBS and pro-iTBS. Our results demonstrate that 100 Hz hdTBS induces after-effects that were three times stronger than those of conventional iTBS.

Since conventional iTBS delivers 600 pulses per session while 100 Hz hdTBS delivers 1200 pulses per session, an interesting question was whether the observed enhancement in TMS effects in the 100 Hz hdTBS group was due to more TMS pulses that these rats had received. To specifically address this question, we have included a pro-iTBS group in our experimental design. pro-iTBS delivers two consecutive iTBS sessions back-to-back, totaling 1200 pulses – matching that of 100 Hz hdTBS. As shown in Fig. 3, in stark contrast to 100 Hz hdTBS, pro-iTBS resulted in a decrease in MEP response. This observation is somewhat counter-intuitive, but consistent with the findings by Gamboe et al., who reported that prolonged iTBS produces an inhibitory effect in the human motor cortex[15].

Taken together, our findings in awake rats, along with Gamboe’s results in humans, underscore the critical roles of the temporal structure of TMS pulses—not merely the total number of pulses—in shaping neuroplasticity. Notably, the pulse pattern of 100 Hz hdTBS most closely resembles the classical TBS protocol and elicited the most pronounced after-effects. Future studies should evaluate the safety and therapeutic potential of 100 Hz hdTBS in humans.

