## Supplemental Figure 1 and Supplemental Table 1 for "High-density theta burst stimulation (hdTBS) at 100 Hz triples the aftereffects of the conventional intermittent TBS"

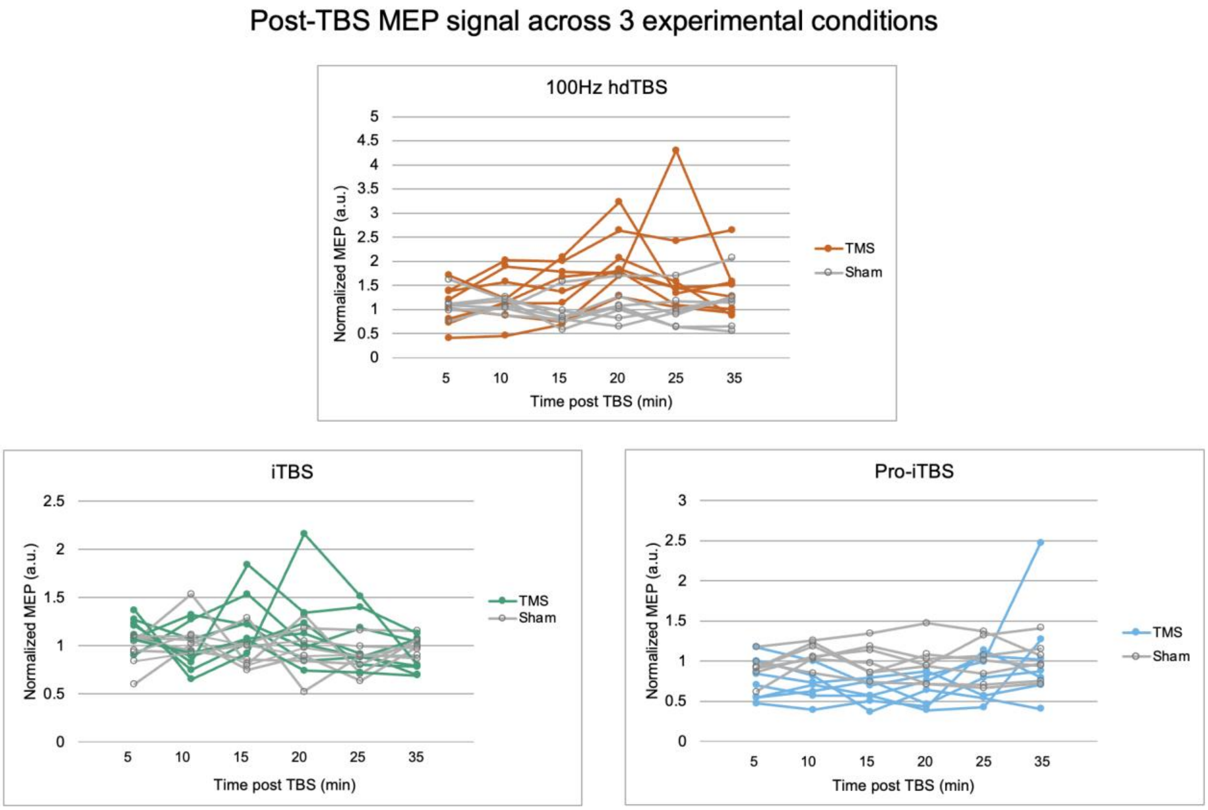

Supplemental Figure 1.

Figure S1. Individual rat MEP measurements at 5, 10, 15, 20, 25, and 35 minutes post-TBS. Each line represents the same rat across each time point, with colors corresponding to treatments received (colored for active TMS, grey for sham).

Supplemental Table 1. Results of post-hoc analysis of AUC values among TBS TYPES

| contrast | estimate | SE | df | t.ratio | p.value | sig. |
|---|---|---|---|---|---|---|
| *100Hz - 100Hz Sham* | 17.75 | 7.041902 | 53 | 2.520626 | 0.045921 | * |
| *100Hz - iTBS* | 17 | 7.041902 | 53 | 2.414121 | 0.046468 | * |
| *100Hz - iTBS Sham* | 29.875 | 7.041902 | 53 | 4.242462 | 0.000832 | *** |
| *100Hz - Prolonged* | 40.21429 | 7.289061 | 53 | 5.517074 | 2.9390410462e-05 | *** |
| *100Hz - Prolonged Sham* | 22.14286 | 7.289061 | 53 | 3.03782 | 0.012927 | * |
| *100Hz Sham - iTBS* | -0.75 | 7.041902 | 53 | -0.10651 | 0.915584 | |
| *100Hz Sham - iTBS Sham* | 12.125 | 7.041902 | 53 | 1.721836 | 0.15913 | |
| *100Hz Sham - Prolonged* | 22.46429 | 7.289061 | 53 | 3.081918 | 0.012927 | * |
| *100Hz Sham - Prolonged Sham* | 4.392857 | 7.289061 | 53 | 0.602664 | 0.591556 | |
| *iTBS - iTBS Sham* | 12.875 | 7.041902 | 53 | 1.828341 | 0.136506 | |
| *iTBS - Prolonged* | 23.21429 | 7.289061 | 53 | 3.184812 | 0.011325 | * |
| *iTBS - Prolonged Sham* | 5.142857 | 7.289061 | 53 | 0.705558 | 0.541579 | |
| *iTBS Sham - Prolonged* | 10.33929 | 7.289061 | 53 | 1.418466 | 0.238602 | |
| *iTBS Sham - Prolonged Sham* | -7.73214 | 7.289061 | 53 | -1.06079 | 0.411036 | |
| *Prolonged - Prolonged Sham* | -18.0714 | 7.52811 | 53 | -2.40053 | 0.046468 | * |
